# A New Minimally Invasive Technique for Repairing Achilles Tendon Rupture:A Biomechanical Study

**DOI:** 10.1101/642520

**Authors:** Peng zhao, Dawei Sun, Yaru Xiong, Ribo Zhuo

## Abstract

**Introduction:** The incidence of Achilles tendon rupture shows a gradually increasing trend, which is mainly managed by minimally invasive treatment due to its advantages, such as low wound infection rate. At present, the firmness of the commonly applied minimally invasive suture method for Achilles tendon remains controversial. Our research group has developed a novel suture method for Achilles tendon, which has achieved favorable clinical outcomes. Therefore, this experiment aimed to explore the optimal approach to repair Achilles tendon rupture through comparing the biomechanical strength of the commonly used Achilles tendon suture methods currently.

**Materials and methods:** 6 fresh frozen human cadaveric Achilles tendon specimens were sutured by three kinds of technique, and were tested through the cyclical loading after repair.

**Results:** Results of cyclical loading showed that, the repair using the new technique was stronger after 10 cycles, 1000 cycles, and rupture. Moreover, the new technique had displayed superior anti-deformation strength to that of the Ma-Griffith technique.

**Conclusions:** Our experimental results demonstrate that, the new technique proposed by our research group can attain comparable biomechanical properties to those of the Krachow technique. However, the sample size in this study is small, and further clinical trials are warranted.

## Introduction

As the strongest tendon in human body, Achilles tendon is of great significance to the movement of the ankle joint, and Achilles tendon rupture can lead to weak plantar flexion ^[1]^. With the continuous improvement in national fitness passion in China, the incidence of acute Achilles tendon injury is increasing year by year^[2]^, which has resulted in the growing requirements for the functional recovery following Achilles tendon repair. Therefore, it is increasingly important to explore a good method to repair the Achilles tendon. In China, the methods to repair Achilles tendon mostly adopt a large incision and will fully expose the tendon broken end after Krachow suture^[3]^. Such method is considered as the “golden standard” for open Achilles tendon repair^[4]^, but such operation has excessively removed the aponeurosis on tendon surface, resulting in great damage to the tendon blood supply. In addition, the Achilles tendon is located at the superficial position, which lacks soft tissue protection, with the postoperative wound infection rate of as high as 21.6%^[5]^. As a result, most scholars have advocated to repair the Achilles tendon through the minimally invasive technique at present^[6]^. In China, the modified Bunnell suture proposed by Ma and Griffith is the commonly adopted minimally invasive suture method^[7]^, however, such method will result in relatively high damage to the sural nerve^[8]^. Some Chinese scholars have reported other minimally invasive surgical methods, such as arthroscopic guidance and ultrasound guidance, nonetheless, they have higher technical and equipment requirements, which are thereby less popular^[9]^. Notably, the cross suture technique proposed by our research group has attained favorable clinical effects on repairing the strength of Achilles tendon and reducing damage to the sural nerve. Thus, we have performed a biomechanical study of 18 cadaveric Achilles tendon to compare the biomechanical characteristics of the cross suture technique with Ma-Griffith, the most commonly used minimally invasive suture method in China, as well as the Krachow suture method that was extensively adopted in China.Our hypothesis was that the cross suture technique would provide a stronger construct than other two techniques.

## Materials and Methods

### Specimen Preparation

In our study, 18 fresh frozen human cadaveric Achilles tendon specimens (male, withthe average age of 35 years, and no injury to the tendon before death) were purchased from the Anatomy Laboratory of Southern Medical University. Each of Achilles tendon specimen was randomly allocated to Ma-Griffith group, Krachow group and the cross suture technique. Before test, the specimens had already been thawed and kept moist by spraying saline.

### Achilles Tendon Repair Method

In this study, the Achilles tendon was cut from the middle, and the Ma-Griffith and Krachow suture techniques for Achilles tendon repair were investigated after standardization. Each group was sutured using the 0.7 mm Ethibond. Fig.1 has showed the detailed cross suture technique.

**Fig.1.**
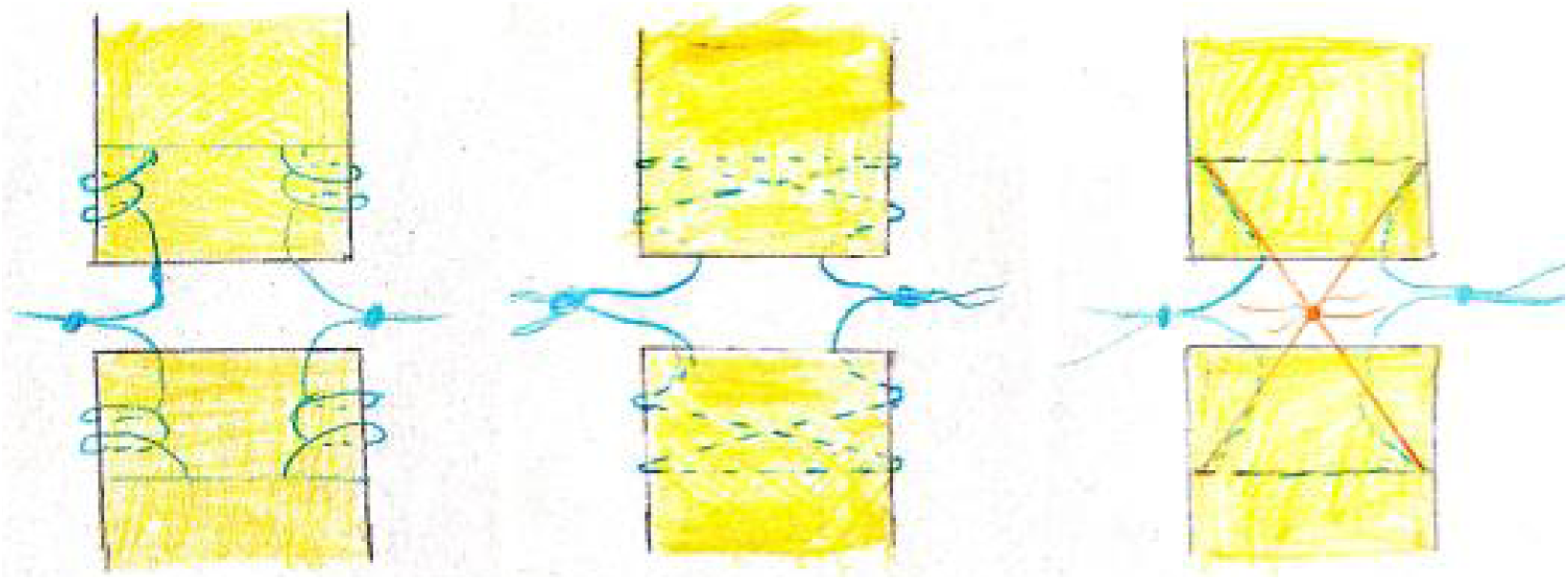
Achilles Tendon Repair Method **A**:Krachow **B**: Ma-Griffith **C**:Cross Suture Technique

### Experimental method

The Achilles tendon in each groups was fixed onto the clamp of the Instron’s Omnipotent Biomechanical Test Machine (ElectroPuls E10000, Instron, USA).Then, the tendon was positioned to be at the center of the clamp, which had been used in various previous studies. Typically, the distance between the two clamps was 4 cm, and a ruler was placed next to the tendon, so as to observe the deformation of the Achilles tendon. Moreover, three stages of test were set. The tension of the machine was set at 30 N-120 N at the first stage, and at 30 N-191 N at the second stage if the Achilles tendon was not ruptured. At the third stage, the tension was increased to 369 N if the Achilles tendon was not ruptured at the second stage. The cycle times of all the three stages were all 1000 times, and the deformation of the Achilles tendon at 10 cycles, 1000 cycles and rupture, together with the number of cycles when it ruptured, would be recorded.

## Statistical analysis

The mean and standard deviation of the measured values would be calculated for all the three groups. The SPSS13.0 software was adopted for all statistical analyses. In addition, ANOVA was applied to evaluate the differences among the three groups (P < 0.05).

## Conclusions

Comparison of the number of cycles among the three groups: the Achilles tendons in the cross suture technique group and the Krachow group had ruptured when entering the third stage after completing 1,000 cycles at the first and second stages, respectively; typically, the 6 samples in Krachow group had ruptured at an average of 2,250 cycles. Besides, the average number of cycles was 2015 in new technology group and 1855 in Ma-Griffith group at the second stage. Meanwhile, difference between the cross suture technique group and Krachow group was not statistically significant (P=1.000), while that between the cross suture technique group and Ma-Griffith group was of statistical significance (P =0.001). In addition, our results suggested that the strength of the new suture technique was comparable to that of Krachow suture, both of which were superior to that of Ma-Griffith suture.

No statistically significant differences in the elongation of the Achilles tendon after 10 cycles [10] were detected among Ma-Griffith group (1.68±0.74) mm, Krachow group (1.79 0.56) mm and new technology group (1.58±0.53) mm (P =0.82), indicating that all the three groups had similar anti-deformation ability after 10 cycles. After the first stage cycle, there was no statistically significant difference in the elongation between the cross suture technique group [(3.11±0.76) mm] and the Krachow group [(3.58±0.58) mm] (P =0.728), but difference between the cross suture technique group [(3.11 0.76) mm] and the Ma-Griffith group [(7.62±1.81) mm] was statistically significant (P =0.000) (table 1). Additionally, difference in the anti-deformation ability between cross suture technique group and Krachow group after 1000 cycles was not statistically significant, and both of them were stronger than that of the Ma-Griffith group.

**Table 1.** Deformation during cycling test **CST(**Cross Suture Technique)

|  | **Elongation after**<br>**10 cycles(mm)** | **Elongation after**<br>**1000cycles(mm)** | **Elongation after**<br>**ruptured(mm)** |
| --- | --- | --- | --- |
| **Ma-Griffith** | **1.68±0.74** | **7.62±1.81** | **8.96±1.7<sup>a</sup>** |
| **Krachow** | **1.79±0.56** | **6.58±0.58** | **6.79±0.88<sup>a</sup>** |
| **CST** | **1.58±0.53** | **6.11±0.76** | **6.51±0.96** |
| **P (P<0.05)** | **0.82** | **0.77** | **0.00** |

<sup>a</sup>p=0.00,vs CST

## Discussion

Compared with the traditional incision and suture, minimally invasive treatment for Achilles tendon rupture is superior in its less damage to the blood supply on the Achilles tendon surface, as well as the remarkably reduced postoperative wound infection rate. Nonetheless, it may cause damage to the sural nerve, and the suture for the Achilles tendon may not be strong enough. As a result, patients are prone to re-rupture during postoperative rehabilitation [11], and thereby required to be immobilized at the ankle joint for a period of time after minimally invasive suture [12], which has induced the occurrence of complications like tendon adhesion. Our research group had set up the three-stage experiment according to the tendon pulling force during rehabilitation. Our experimental results showed that for the Ma-Griffith suture technique, the Achilles tendon ruptured at the second stage of test, while for the cross suture technique of suture that we proposed, the Achilles tendon could smoothly experience the third stage of test, which ruptured in the 2015th cycle, while the Achilles tendon had ruptured in the 2250th cycle for the Krachow suture rupture technique at the third stage. Obviously, the strong degree of the new technique was superior to that of the Ma-Griffith suture technique. Therefore, the cross suture technique that we proposed could provide stabilized fixation strength for the early rehabilitation training of patients.

It was found when observing the rupture Achilles tendon end and the tendon deformation that, the Krachow and Ma-Griffith sutures would split at the suture knot, since the single stitches tendon knot in place could easily produce stress concentration. When tension was applied to the tendon repaired by the cross suture technique, the parallelogram-like deformation occurred in the muscle, which could convert the stress generated by the thread knot into the pressure onto the Achilles tendon broken end, reduce the tendon ectropion resulted from the tightening of the tendon suture, increase the tendon contact area, and promote Achilles tendon healing.

The rupture and repair of the Achilles tendon will frequently cause weak heel lift due to tendon extension after suturing[13], which would bring about great psychological burdens on the patients. In this experiment, after 1000 cycles and the measurement of the ruptured posterior Achilles tendon, no significant difference was found in the elongation of the three suture methods within 1000 cycles, but the Krachow suture method had the shortest elongation at the time of rupture, while that of the cross suture technique was shorter than that of the Ma-Griffith suture method but was still longer than that of the Krachow suture method. Therefore, the cross suture technique could effectively prevent the occurrence of tendon elongation.

In this experiment, percutaneous suture channels were established to keep the sural nerve out of the suture channel, thus avoiding injury to the sural nerve. However, large clinical trial support is lacking in this study, as a result, further research and exploration are needed.

## References

[1] Padanilam, Thomas G. Chronic Achilles Tendon Ruptures[J]. Foot & Ankle Clinics of North America, 2009, 14(4):711–728.

[2] Majewski M, Rickert M, Steinbrück, K. [Achilles tendon rupture. A prospective study assessing various treatment possibilities][J]. Der Orthopde, 2000, 29(7):670–6.

[3] Clement D B, Taunton J E, Smart G W. Achilles tendinitis and peritendinitis: Etiology and treatment[J]. The American Journal of Sports Medicine, 1984, 12(3):179–184.

[4] Nonoperative, Dynamic Treatment of Acute Achilles Tendon Rupture: Influence of Early Weightbearing on Biomechanical Properties of the Plantar Flexor Muscle–Tendon Complex—A Blinded, Randomized, Controlled Trial[J]. The Journal of Foot and Ankle Surgery, 2015, 54(2):220–226.

[5] Kocher M S, Bishop J, Marshall R, et al. Operative versus nonoperative management of acute Achilles tendon rupture - Expected-value decision analysis[J]. American Journal of Sports Medicine, 2002, 30(6):783.

[6] Kadakia A R, Dekker R G, Ho B S. Acute Achilles Tendon Ruptures: An Update on Treatment[J]. Journal of the American Academy of Orthopaedic Surgeons, 2017, 25(1):23.

[7] Ma G W C, Griffith T G. Percutaneous Repair of Acute Closed Ruptured Achilles Tendon[J]. Clinical Orthopaedics and Related Research, 1977, &NA;(128):247–255.

[8] Aibinder W R, Patel A, Arnouk J, et al. The Rate of Sural Nerve Violation Using the Achillon Device: A Cadaveric Study[J]. Foot & Ankle International, 2013, 34(6):870–875.

[9] Lacoste S, J M. Féron, Cherrier B,. Percutaneous Tenolig(?) repair under intra-operative ultrasonography guidance in acute Achilles tendon rupture[J]. Orthopaedics & Traumatology Surgery & Research Otsr, 2014, 100(8):925–30.

[10] Lacoste S, Féron J M, Cherrier B. Percutaneous Tenolig(®) repair under intra-operative ultrasonography guidance in acute Achilles tendon rupture.[J]. Orthopaedics & Traumatology Surgery & Research Otsr, 2014, 100(8):925–30.

[11] Mcquillan R, Gregan P. Tendon rupture as a complication of corticosteroid therapy [2][J]. Palliative Medicine, 2005, 19(4):352–3.

[12] Demetracopoulos C A, Gilbert S L, Young E, et al. Limited-Open Achilles Tendon Repair Using Locking Sutures Versus Nonlocking Sutures: An In Vitro Model[J]. Foot & ankle international. / American Orthopaedic Foot and Ankle Society [and] Swiss Foot and Ankle Society, 2014, 35(6):612–618.

[13] Padanilam, Thomas G. Chronic Achilles Tendon Ruptures[J]. Foot & Ankle Clinics of North America, 2009, 14(4):711–728.

